# User-Assisted Approach for Accurate Nonrigid Registration of Images and Traces

**DOI:** 10.1101/2025.01.29.635549

**Authors:** Mirza M. Junaid Baig, Tina Thuy N. Hoang, Samuel H. Chung, Armen Stepanyants

## Abstract

Fully automated registration algorithms are prone to getting trapped in solutions corresponding to local minima of their objective functions, leading to errors that are easy to detect but challenging to correct. Traditional solutions often involve iterative parameter tuning, data preprocessing and preregistering, and multiple algorithm reruns—an approach that is both time-consuming and does not guarantee satisfactory results. Therefore, for tasks where registration accuracy is more important than speed, it is appropriate to explore alternative, user-assisted registration strategies. In such tasks, finding and correcting errors in automated registration is often more time-consuming than directly integrating user input during the registration process.

Therefore, this study evaluates a user-assisted approach for accurate nonrigid registration of images and traces. By leveraging the corresponding sets of fiducial points provided by the user to guide the registration, the algorithm computes an optimal nonrigid transformation that combines linear and nonlinear components. Our findings demonstrate that the registration accuracy of this approach improves consistently with the increased complexity of the linear transformation and as more fiducial points are provided. As a result, accuracy sufficient for many biomedical applications can be achieved within minutes, requiring only a small number of user-provided fiducial points.

## 1. Introduction

Registration is a fundamental tool for computer vision [1, 2], biological [3], and medical [4, 5] applications. It enables quantitative comparisons and meaningful combinations of datasets required for the detection of changes in longitudinal studies [6, 7], cross-subject comparisons [8], and multimodal data integration [9, 10]. Despite its broad applicability, registration remains challenging, particularly in a nonrigid context. Fully automated algorithms offer efficiency but can struggle with large rotations and deformations, high structural variability, and experimental artifacts [11]. Such methods usually rely on simplistic models and can get trapped in local minima leading to erroneous solutions. For instance, even state-of-the-art fully automated algorithms for point set registration such as Coherent Point Drift (CPD) [12, 13] and Bayesian Coherent Point Drift (BCPD) [14] are prone to such problems (Figure 1A). Preregistering the data can improve the outcome but often falls short of achieving satisfactory results (Figure 1B). Extensive parameter tuning may improve performance but undermines the goal of full automation.

**Figure 1.**
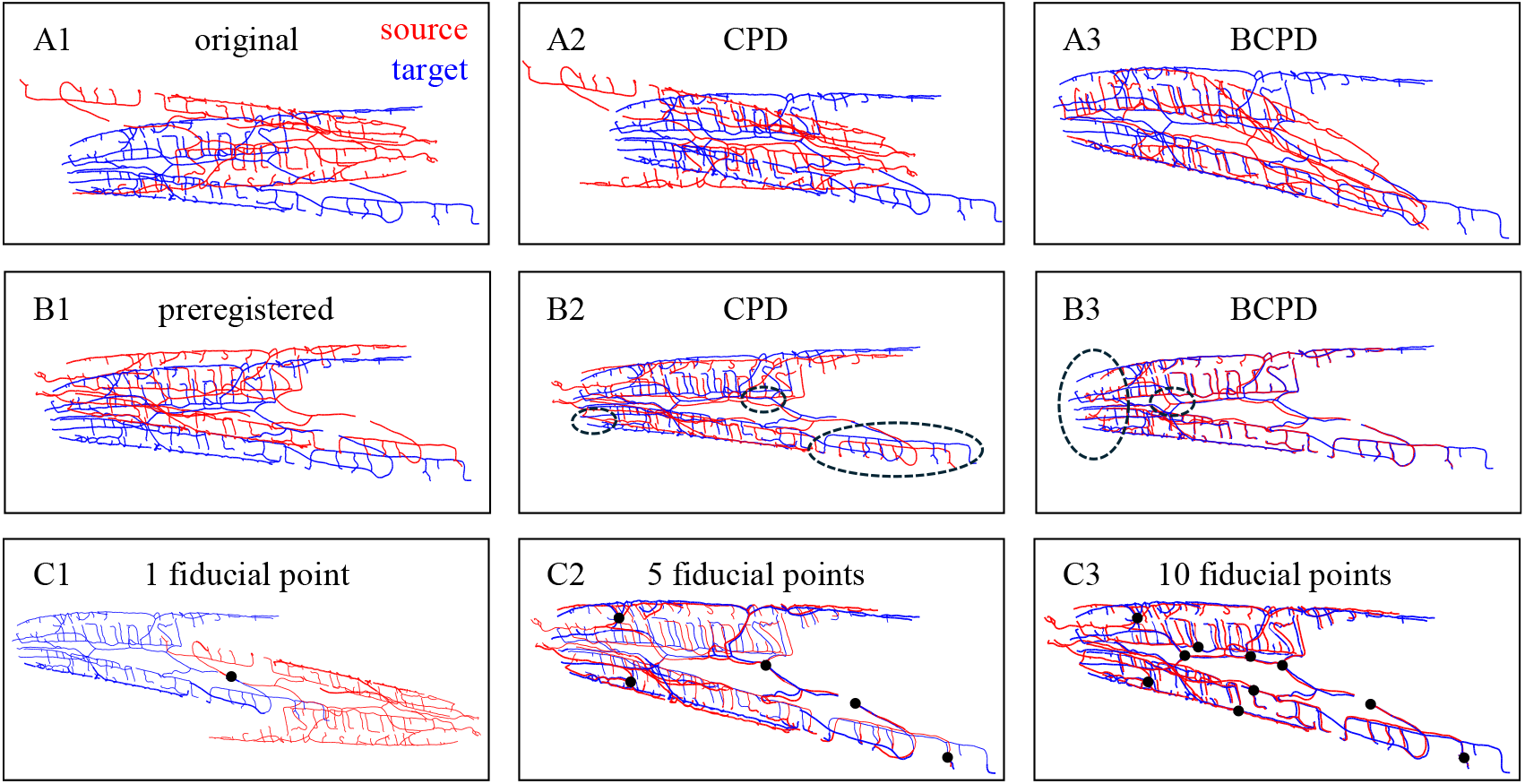
User-assisted registration circumvents problems associated with fully automated algorithms. **A.** Fully automated registration often converges to suboptimal solutions. This is illustrated through the registration of 3D traces of FLP neurons using CPD and BCPD algorithms with default parameters. **B**. After the user preregisters the traces, performance improves but does not yield satisfactory results (circled regions). **C**. User-assisted registration gradually improves with an increasing number of user-defined fiducial points, leading to a superior result.

Instead, user-assisted methods can provide more efficient and reliable alternatives from the start. In such methods, expert users provide key corresponding landmarks, addressing many challenges of full automation by prioritizing accuracy over speed. The accuracy of user-assisted methods depends on user experience and effort (Figure 1C), and the desired outcome can be achieved in any application given a suitable registration model. Therefore, such methods can yield superior performance, making them well-suited for complex biomedical applications.

Many recent advances in registration have been driven by Deep Learning (DL) [15]. Techniques such as Reinforcement Learning [16, 17] and Generative Adversarial Networks [16, 18, 19] have demonstrated impressive improvements in accuracy and robustness. The power of DL-based methods lies in their ability to generalize across datasets and achieve good results without direct user involvement once the model is created. However, DL-based methods require extensive training datasets which are arduous to create and incur high computational costs. In addition, their reliance on black-box models that lack interpretability makes it difficult to address domain-specific constraints to ensure meaningful outcomes [16, 20]. Thus, while DL-based methods have a significant potential for advancing the field of registration, at present, they are not the best fit for biomedical applications where accuracy, interpretability, and flexibility are paramount.

The user-assisted registration approach described in this study offers a complementary solution that combines user expertise with algorithmic efficiency, enabling high accuracy while maintaining interpretability. The registration problem is formulated as an exactly solvable model in which linear and nonlinear transformations are combined to achieve the exact registration of user-defined fiducial points. Four types of linear transformations are considered, including translation, rigid, similarity, and affine. Additionally, a custom treatment of similarity and affine transformations is introduced in which an optimal rotation is discounted from their associated costs. This makes it possible to test the idea that increasing the complexity of linear transformation minimizes the warping of the space.

The proposed method supports the registration of 2D and 3D point sets, traces, and images. It is equipped with a graphical interface, allowing users to visually select corresponding points and iteratively improve the registration outcome (Figure 1C). This approach is particularly useful for domain scientists working with specialized datasets for which there are no immediate registration solutions available. By leveraging user expertise and focusing on interpretability, this approach provides a reliable alternative to DL-based methods, particularly in applications demanding domain-specific precision and flexibility. We demonstrate the utility of this approach on the registration of 3D traces of *C. elegans* neurons and 2D images of the human retina, showing significant improvements in registration accuracy over fully automated methods.

## 2. Model for exact registration of fiducial points with a combination of linear and nonlinear transformations

Consider point sets *X*, referred to as the source, and *Y*, referred to as the target, in ℝ^*D*^ containing *N* user-defined fiducial points each (Figure 2A). Positions of the points are denoted with vectors **x**_*i*_ and **y**_*i*_ respectively (*i* = 1, …, *N*), and their correspondences are known. The point set *X* must be transformed and registered to *Y*, which remains fixed. The registering transformation, **T**(**x**) = **Ax** + **b** + **V**(**x**), which consists of a linear component governed by matrix **A** and vector **b**, and a nonlinear part described by a vector-valued function **V**(**x**), must match the corresponding fiducial points exactly, i.e., **T**(**x**_*i*_) = **Ax**_*i*_ + **b** + **V**(**x**_*i*_) for all *i* (Figure 2A). The nonlinear part of the registering transformation is modeled with the Coherent Point Drift (CPD) method of Myronenko et al. [13], and four types of linear transformations of increasing complexity are considered:

**Figure 2.**
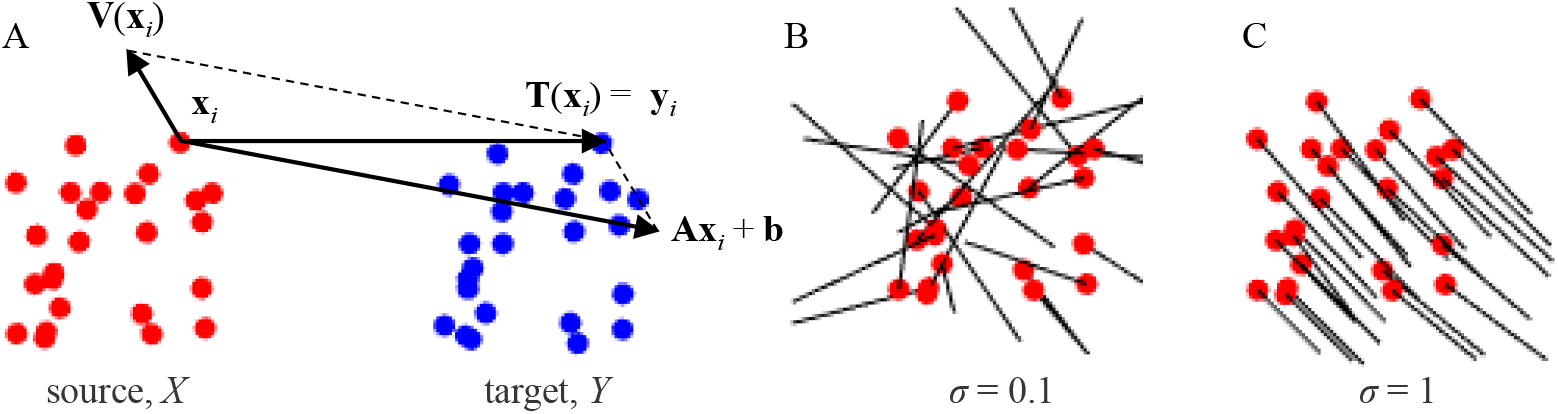
Registering transformation. **A.**Source and target point sets with known correspondences before registration. The sum of displacement vectors for linear and nonlinear parts of the registering transformation for each source point (**x**_*i*_) results in a displacement that exactly matches the position of the corresponding target point (**y**_*i*_). **B**. Nonlinear displacement field of the source points, **V**(**x**_*i*_), is indicated with black lines. **C**. Coherence of the nonlinear displacement field increases with *σ*. Point set *X* was randomly drawn from a unit cube and *Y* was generated from *X* by adding random displacements in the [−0.1 0.1] range. Registration was performed with an affine + nonlinear transformation with *λ* = 0.

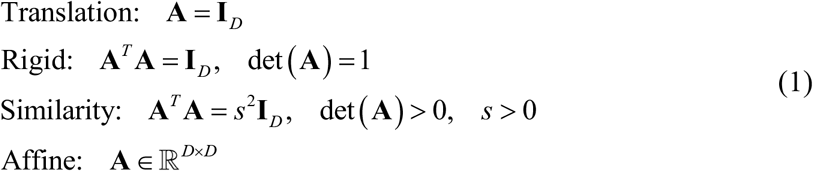

In these expressions and below, bold lower-case letters represent *D*×1 vectors, bold upper-case characters denote *D*×*D* matrices or, in the cases of **T**(**x**) and **V**(**x**), *D*×1 vector-valued functions. The trace of a matrix is denoted as tr and **I**_*D*_ represents a *D*×*D* identity matrix. In the similarity case, **A** can be written as a product of a uniform scaling factor *s* and a rotation matrix.

Parameters of the registering transformation are determined through the minimization of the heuristic cost of space deformation associated with the nonrigid nature of **T**(**x**), subject to constraints on this transformation.

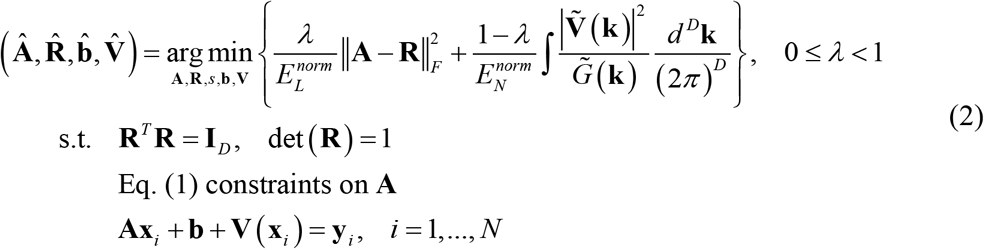

The heuristic cost of Eq. (2) is a weighted sum of normalized costs associated with the linear (*E_L_*) and nonlinear (*E_N_*) deformations induced by **T**(**x**). Parameter *λ* shifts the balance of these contributions from a purely nonlinear cost at *λ* = 0 to a purely linear cost in the *λ* → 1 limit. The linear cost component is a normalized squared Frobenius norm of the difference between **A** and an orthogonal matrix with unit determinant, **R** (rotation without reflection). The latter matrix is determined during optimization and ensures that an optimal rotation is discounted from the linear deformation cost. The nonlinear cost component is a normalized CPD metric [13], where 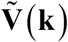 represents a Fourier transform of **V(x)** and 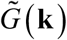 denotes a Fourier transform of a Gaussian function.

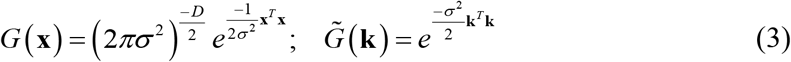

Parameter *σ* is referred to as the coherence parameter, for the reason that the structure of the nonlinear cost component forces the source points that are within distance *σ* of each other to move coherently (in similar directions) during the transformation [13] (Figure 2B, C).

The normalization factors for the linear and nonlinear cost components,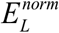 and 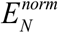 in Eq. (2) are included to ensure that the two cost terms have similar scales, making it easier to interpret the tradeoffs of their contributions. These normalization factors will be deduced following the solution of the optimization problem and will depend on the positions of the source and target points and the coherence parameter, but not on *λ* and the type of linear transformation. This will enable comparisons of registration results when different transformations are applied to the same point sets.

The described registration model is governed by only two parameters, *λ* and *σ*, and the outcome of registration depends on the choice of the linear transformation and fiducial points provided by the user.

## 3. Solution of the model

The heuristic cost of Eq. (2) must be minimized with respect to variables **A, R**, *s* (similarity only), **b**, and function **V**(**x**), subject to constraints. This is done gradually starting with the optimization of function **V**. To that end, the third set of constraints of Eq. (2) is included in the objective function using Lagrange multipliers **c**_*i*_, and the optimization is performed with standard methods of variational calculus.

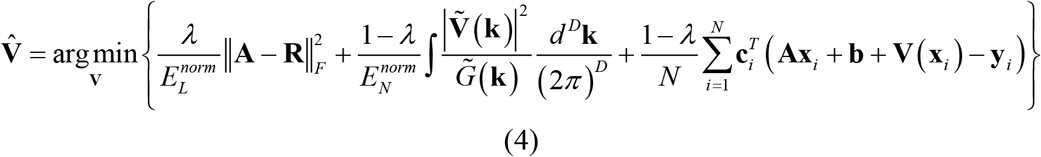

The result shows that 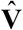 is a mixture of Gaussians, and the Lagrange multipliers **c**_*i*_ depend on the inverse of the Gaussian kernel, *𝒢*= {*G* **(x**_*i*_ − **x**_*j*_)}∈ℝ ^*N* × *N*^.

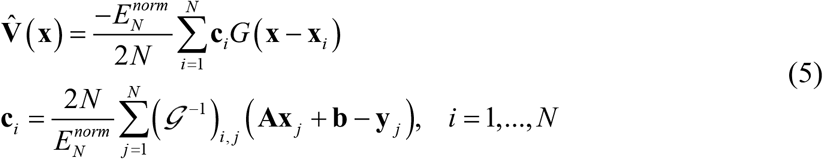

Next, after eliminating 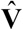 from the original optimization problem of Eq. (2), the process is continued by performing minimization with respect to **b**.

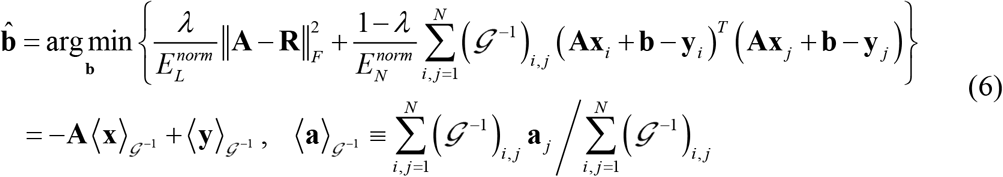

The remaining problem can be concisely written with the help of modified covariance matrices 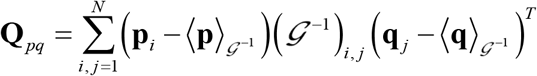.

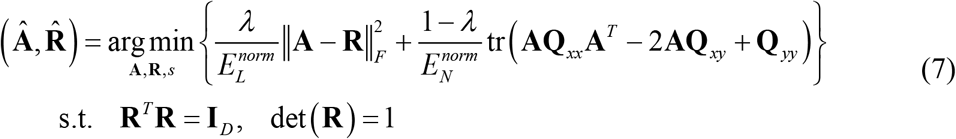

This problem is solved by employing the singular value decomposition (SVD) trick [21], which helps to enforce the constraints on matrix **R**. The results for the four types of linear transformations are summarized in Table 1.

**Table 1.**
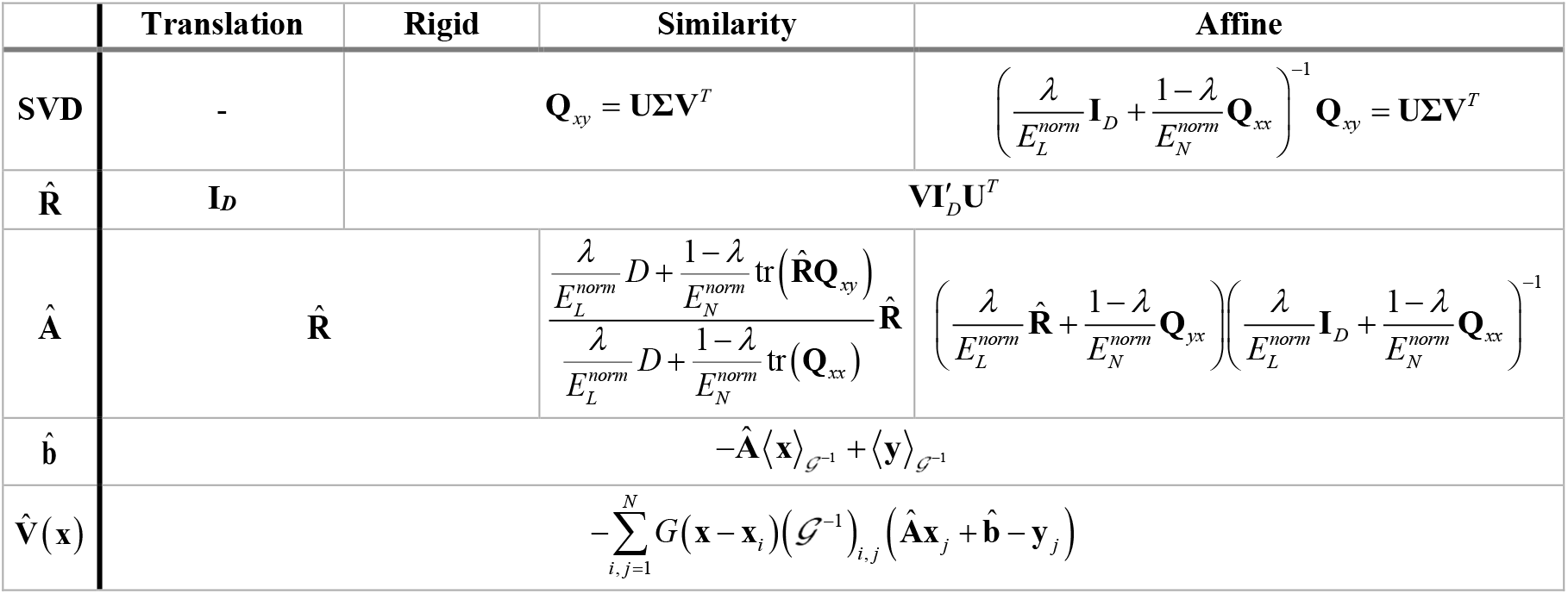
Model solutions for the considered linear transformation types.

The solution for each type of linear transformation must be constructed by executing the rows of Table 1 from top to bottom. Matrix 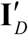 which appears in the expressions for 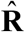 is the identity matrix **I**_*D*_ with 1 in the last row and column replaced with det **(VU**^*T*^ ). The nonlinear part of the registering transformation ensures that, so long as points in set *X* are in general positions, any registration problem is feasible, i.e. an optimal solution satisfying the constraints of Eq. (2) can be found. The coherence of the nonlinear displacement field, **V**(**x**), is governed by parameter *σ*, such that points within *σ* move in similar directions, Eq. (5) (Figure 2B, C).

The normalized costs of linear and nonlinear deformations, *E_L_* and *E_N_*, are calculated based on the optimal transformation.

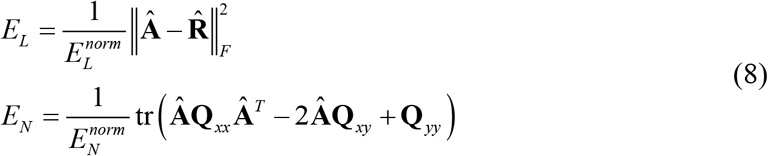

It remains to determine the normalization factors 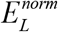 and 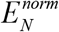 to ensure that, for the given *σ* and point sets, *E_L_* and *E_N_* as functions of *λ* have similar bounds for all linear transformation types.

To accomplish this, we examine the dependence of these cost components on *λ*. When *λ* = 0, the burden of registration is placed maximally on the linear transformation because it incurs no penalty. Therefore, the maximal *E_L_* is expected at *λ* = 0 for all linear transformation types. Similarly, in the *λ* → 1 limit, the nonlinear part of the transformation does not contribute to the heuristic cost of Eq. (2) and is expected to cause the largest spatial distortion resulting in a maximal *E_N_*. This reasoning is confirmed by direct analytical calculations, and an example of the dependence of the normalized cost components on *λ* is shown in Figure 3C. Therefore, we focus on these limiting cases and show the explicit expressions for *E_L_* at *λ* = 0 and *E_N_* at *λ* → 1 for all linear transformation types (Table 2).

**Table 2.** Expressions for *E_L_* at *λ* = 0 and *E_N_* at *λ* → 1 for the optimal transformations from Table 1.

|  |  | Translation | Rigid | Similarity | Affine |
| --- | --- | --- | --- | --- | --- |
| $\lambda = 0$ | SVD | - | $\mathbf{Q}_{xy} = \mathbf{U}\Sigma\mathbf{V}^T$ | | $\mathbf{Q}_{xx}^{-1}\mathbf{Q}_{xy} = \mathbf{U}\Sigma\mathbf{V}^T$ |
| | $E_L$ | 0 | | $\frac{D}{E_L^{norm}} \left( 1 - \frac{\text{tr}(\mathbf{I}'_D \Sigma)}{\text{tr}(\mathbf{Q}_{xx})} \right)^2$ | $\frac{\text{tr}((\mathbf{I}'_D - \Sigma)^2)}{E_L^{norm}}$ |
| $\lambda \rightarrow 1$ | SVD | - | $\mathbf{Q}_{xy} = \mathbf{U}\Sigma\mathbf{V}^T$ | | |
| | $E_N$ | $\frac{\text{tr}(\mathbf{Q}_{xx} - 2\mathbf{Q}_{xy} + \mathbf{Q}_{yy})}{E_N^{norm}}$ | $\frac{\text{tr}(\mathbf{Q}_{xx} - 2\mathbf{I}'_D \Sigma + \mathbf{Q}_{yy})}{E_N^{norm}}$ | | |

**Figure 3:**
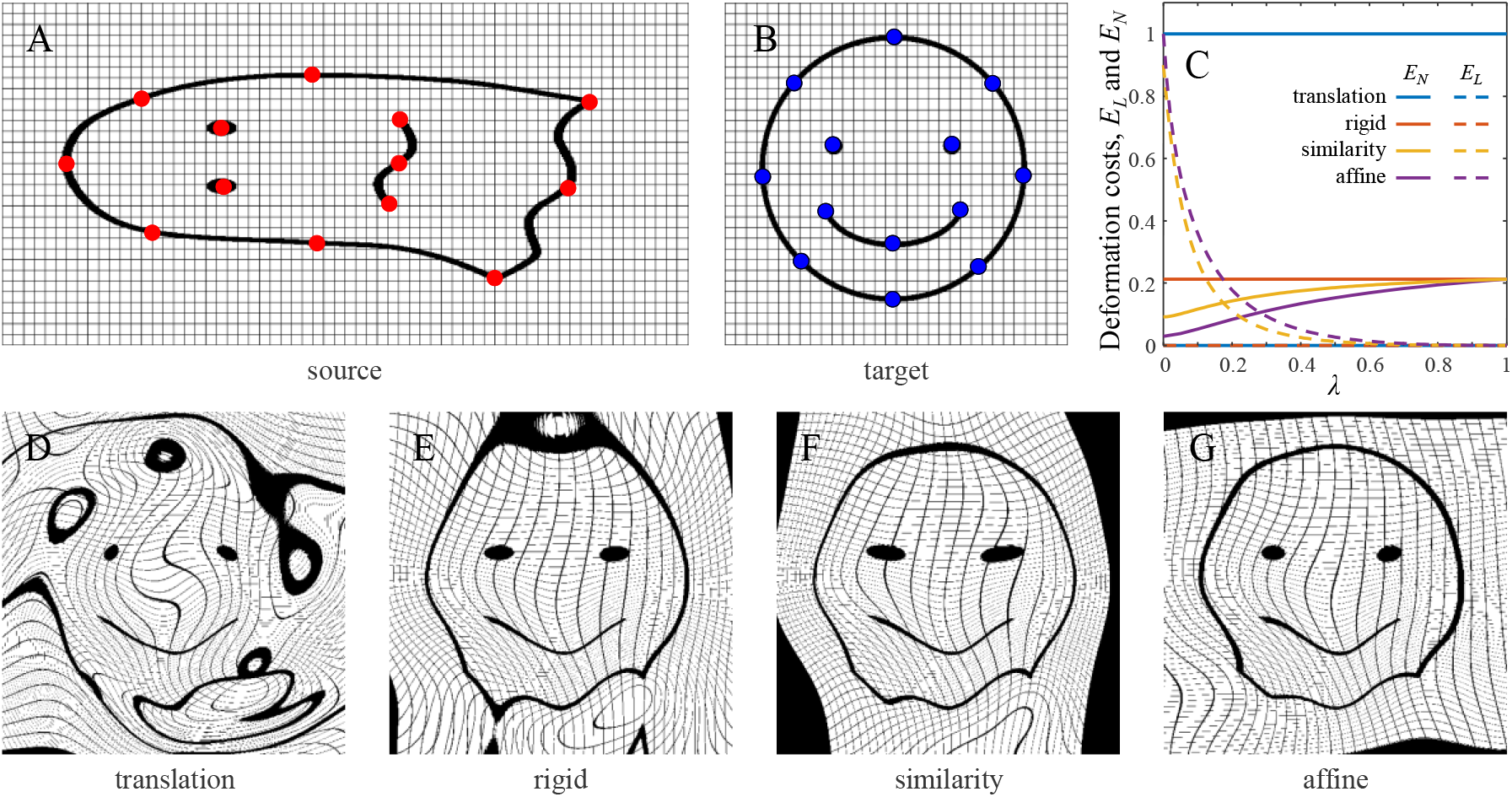
Increasing the complexity of linear transformation reduces space distortion resulting from registration. **A**: Sample source image with 13 fiducial points marked in red (513×1026 pixels). **B**: Target image with the corresponding fiducial points marked in blue (513×513 pixels). **C.** Normalized deformation costs associated with the linear (*E_L_*) and nonlinear (*E_N_*) parts of the registering transformation as functions of parameter *λ* (*σ* = 75). **D-G**. Results of registration for the considered transformation types. The fiducial points are registered exactly by all transformations, and the warping of the space decreases systematically from translation to rigid to similarity to affine (*σ* = 75 and *λ* = 0).

Because the affine transformation is more flexible than the rest, it usually incurs a higher cost (see Figure 3C for an example).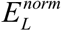 is chosen to make the normalized cost of this transformation equal to 1 at *λ* = 0. On the other hand, because translation is the simplest of the considered transformations, it is accompanied by the largest nonlinear deformation, and 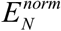 is set in a way that makes the normalized cost of this deformation equal to 1.

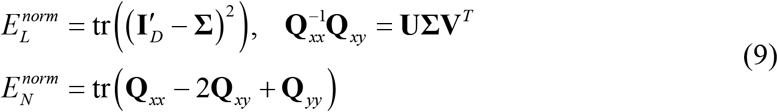

## 4. Linear transformation is necessary to minimize warping of the space

In many biomedical applications, the bulk of the registration task can be solved by a linear transformation, requiring only a small nonlinear correction. In such cases, explicitly including both contributions into a registering transformation as done in this study, **T**(**x**) = **b** + **Ax** + **V**(**x**), can help minimize the warping of the space associated with **V**(**x**). This formulation can be thought of as the Taylor series expansion of **T**(**x**) to the first order in **x**, in which **V**(**x**) corresponds to the remainder term.

To illustrate this idea, fiducial points selected on a synthetic source and target image pair were registered with different parameter settings (Figure 3A, B), and the resulting optimal transformations were applied to the source image. The registration framework guarantees that the fiducial points register exactly, however, the type of linear transformation and parameters *λ* and *σ* can influence the warping of the space. This was confirmed by analyzing the normalized deformation costs associated with linear and nonlinear parts of the registering transformations (Figure 3C) and visually examining the registered images (Figure 3D-G). For translation and rigid transformations, which incur no linear penalty, *E_L_* and *E_N_* are independent of *λ* since the first term in the heuristic cost of Eq. (2) becomes zero. For similarity and affine, there is a tradeoff between *E_L_* and *E_N_*, indicating that these linear transformations help reduce the cost of nonlinear distortion. Importantly, Figure 3C shows that increasing the complexity of linear transformation results in a lower *E_N_*, suggesting that this can help reduce space distortion. This is confirmed in Figure 3D-G which illustrates a progressive reduction in distortion of the grid from translation to rigid to similarity to affine.

## 5. Registration of traces and images

To test the performance of the registration algorithm on biomedical data, the method was applied to a custom dataset of neuron traces and a standard dataset of retinal images.

The neuron traces were obtained from young adult hermaphrodite *C. elegans*, in which FLP neurons were fluorescently labeled with (green fluorescent protein) GFP [22] and imaged *in vivo* in 3D with a confocal microscope. Each image stack consisted of 2048 × 2048 pixels (203 × 203 µm) in the *x* and *y* dimensions and 40 *z*-planes (1 µm *z*-step). The images were traced semi-automatically in 3D using NCTracer software [23, 24] (Figure 4A1, 2). A neuron trace captures the primary structure of a neuron as a series of points connected with line segments. Each *C. elegans* was imaged twice within 20 minutes during which time the animal was moved to a new slide to introduce nonlinear deformations. The close temporal proximity of the imaging sessions ensured minimal structural changes in neuron morphology. This process was repeated for three individual *C. elegans*, yielding three pairs of neuron traces for registration testing.

**Figure 4.**
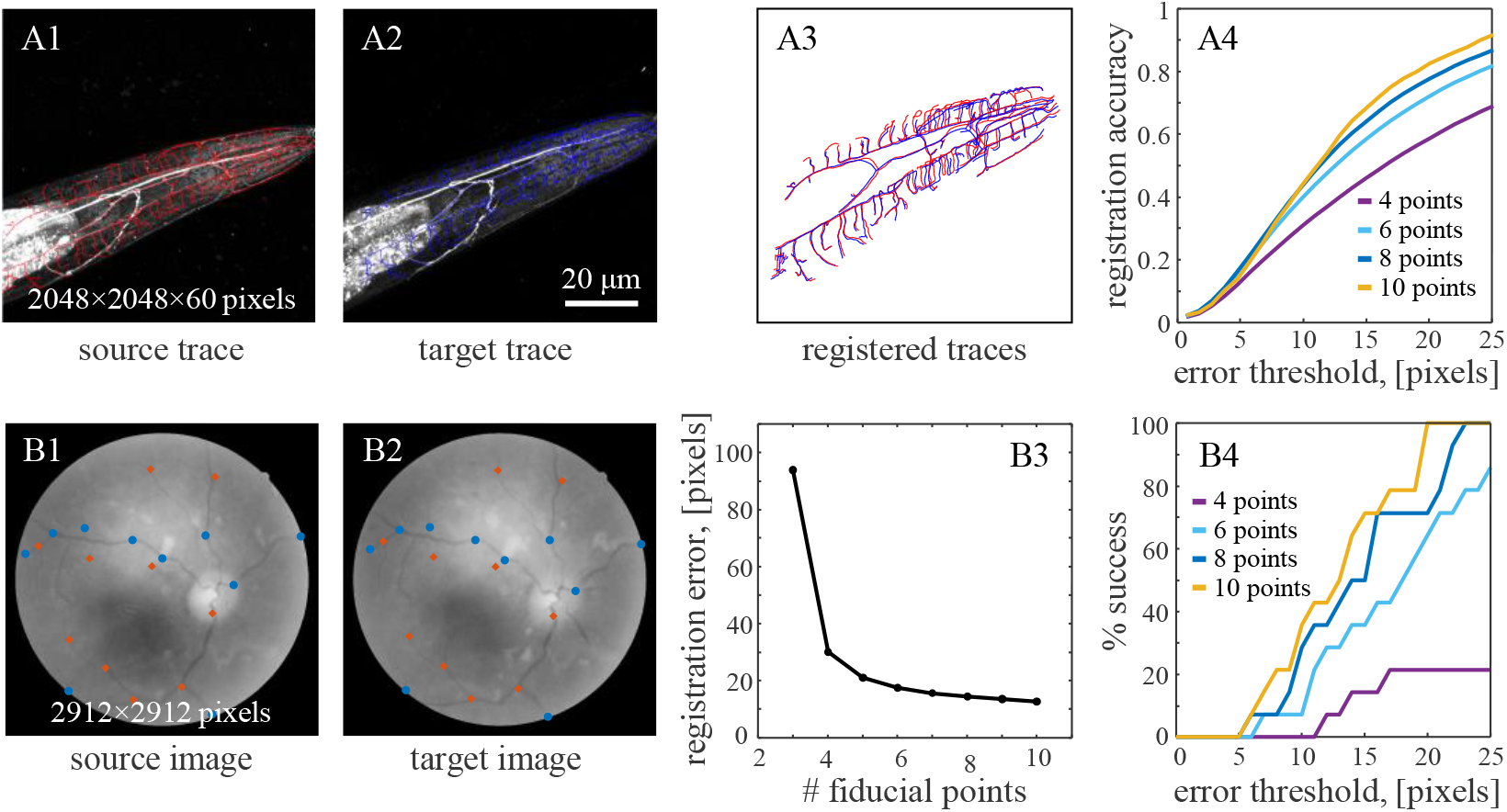
Applications of the user-assisted registration method to biomedical problems. **A1.** Maximum intensity projection of a 3D image stack of an FLP neuron in *C. elegans*. The trace of its axon and dendrites (red) is used as a source for registration. **A2**. Same for the target image and trace. **A3**. Registered source and target traces from (A1) and (A2) using 10 fiducial points (affine + nonlinear, *σ* = 100, *λ* = 0.5). **A4**. Registration accuracy as a function of error threshold systematically improves with an increasing number of user-provided fiducial points (*n* = 3 neuron pairs). **B1, 2**. Sample source and target images from Category A of the FIRE dataset of retinal images, showing ground truth points (red) and user-selected fiducial points (blue). **B3**. Registration error decreases with an increasing number of fiducial points (*n* = 14 image pairs, affine + nonlinear, *σ* = 100, *λ* = 0.5). **B4**. The percentage of successfully registered image pairs as a function of the error threshold systematically increases with the number of fiducial points.

To facilitate registration, a MATLAB graphical user interface (GUI) was developed for selecting corresponding fiducial points in source and target data. The user-selected fiducial points served as inputs to the registration algorithm, and the output transformation was subsequently applied to the entire source trace (Figure 4A3). Registration was performed using an affine + nonlinear transformation and its accuracy was assessed by calculating the fraction of corresponding trace points between the target and registered source traces. Here, two points were deemed corresponding if their Euclidean distance was below a given threshold. Figure 4A4 shows the dependence of registration accuracy on the number of fiducial points and the threshold. It illustrates that accuracy gradually improves with the number of fiducial points, and > 0.9 accuracy is achieved at an error threshold of 25 pixels with only 10 fiducial points.

Next, the performance of the registration method was tested on the Fundus Image Registration (FIRE) dataset [25] containing pairs of human retinal images acquired at different examinations. This dataset provides ground truth points for each image pair, allowing for quantitative evaluation of registration performance. Specifically, Category A of this dataset was examined because it features 14 pairs of images containing anatomical differences due to the progression or remission of retinopathy, making it the most challenging category for registration [26]. Fiducial points for registration were selected using the GUI without prior knowledge of the ground truth points to avoid bias, and selected points within 100 pixels of a ground truth point were removed (Figure 4B1,2).

Registration was performed using an affine + nonlinear transformation and its error was assessed by calculating the average Euclidean distance between the corresponding ground truth points on the target and registered source images. Figure 4B3 shows that this registration error decreases as more fiducial points are used for registration, and an error of < 13 pixels is attained with 10 fiducial point pairs. In addition, the standard registration success rate [25] was calculated as the percentage of registered source ground truth points that were within an error threshold of their corresponding target ground truth points. Figure 4B4 shows that the user-assisted algorithm achieves a 100% registration success rate at an error threshold of 20 pixels with only 10 fiducial point pairs. This result favorably compares to the best fully automated methods [26-29].

## 6. Discussion

This study demonstrates that user-assisted registration can be accurate and efficient. By incorporating fiducial points selected by the user, the method reduces the need for iterative parameter tuning associated with fully automated algorithms, making it a practical choice for real-world applications. The versatility of this approach extends across a range of data modalities and applications. The data may involve 2D and 3D point sets, traces, and images, while the applications may include common biomedical tasks such as registration of neural traces, vascular structures, labeled tissues, and CT scans. For instance, this method can provide precise alignment of neuron traces, enabling the quantitative comparisons needed for *in vivo* longitudinal studies of structural changes associated with learning and memory formation [30-32].

Accuracy of the described registration method systematically improves with the number of fiducial points (Figure 4) and may benefit from using a less constrained linear transformation such as similarity or affine to reduce warping of the space (Figure 3). However, we find that using a linear transformation that matches the registration task is the best practice for achieving robust results. In combining user input with automation, this method has the potential to achieve a balance between accuracy and efficiency required in many registration tasks.

The registration GUI, sample images, and the dataset of neural traces used in this study are available on the Neurogeometry group GitHub site, https://github.com/neurogeometry/User-Assisted-Registration.

## Acknowledgements

This work was supported by the NIH grant R56 NS128413.

